## Supplementary Information for "Single-cell eQTL mapping in yeast reveals a tradeoff between growth and reproduction"

### Supplementary Figures

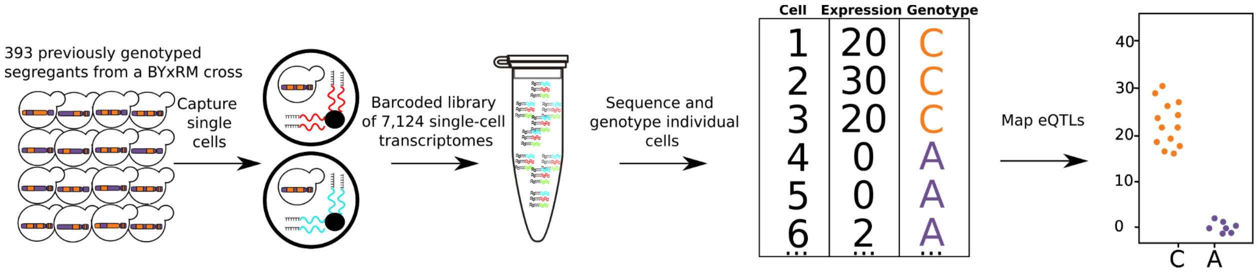


**Figure S1: Single-cell eQTL mapping of 393 previously genotyped segregants.** 393 segregants were pooled, grown in minimal medium, and processed using the 10x Chromium device. The resulting barcoded library is sequenced with Illumina short-read sequencing. The number of supporting molecules for each allele is inferred at every variant position between the parental strains and a hidden Markov model is used to infer the genotype of each segregant. A cartoon example of one eQTL is shown on the top right, cells with the C allele had higher expression than cells with A allele.

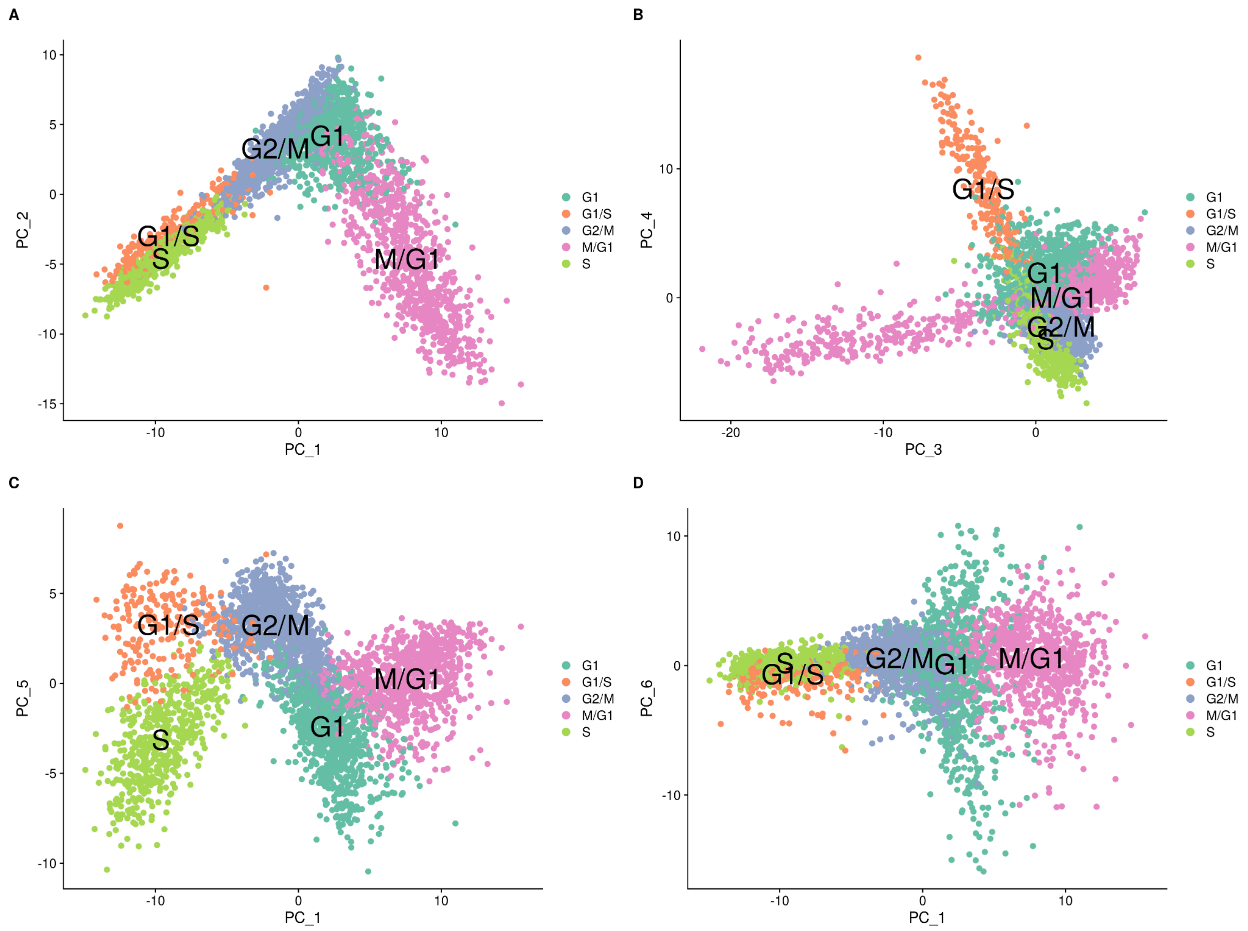


**Figure S2: Cell-cycle classification of the single-cells from the set of 393 previously genotyped segregants visualized on different combinations of principal components.** These data are from the first 10x run of the 393 segregants (Number of cells=3,454). The results look similar for the second run. The principal components (PCs) were calculated using Seurat with the cell-cycle genes. Cells are colored according to their assigned cell cycle stage. **A)** PC plot from the single-cell data comparing PC 1 and PC 2. **B)** PC plot from the single-cell data comparing PC 3 and PC 4. **C)** PC plot from the single-cell data comparing PC 1 and PC 5. **D)** PC plot from the single-cell data comparing PC 1 and PC 6.

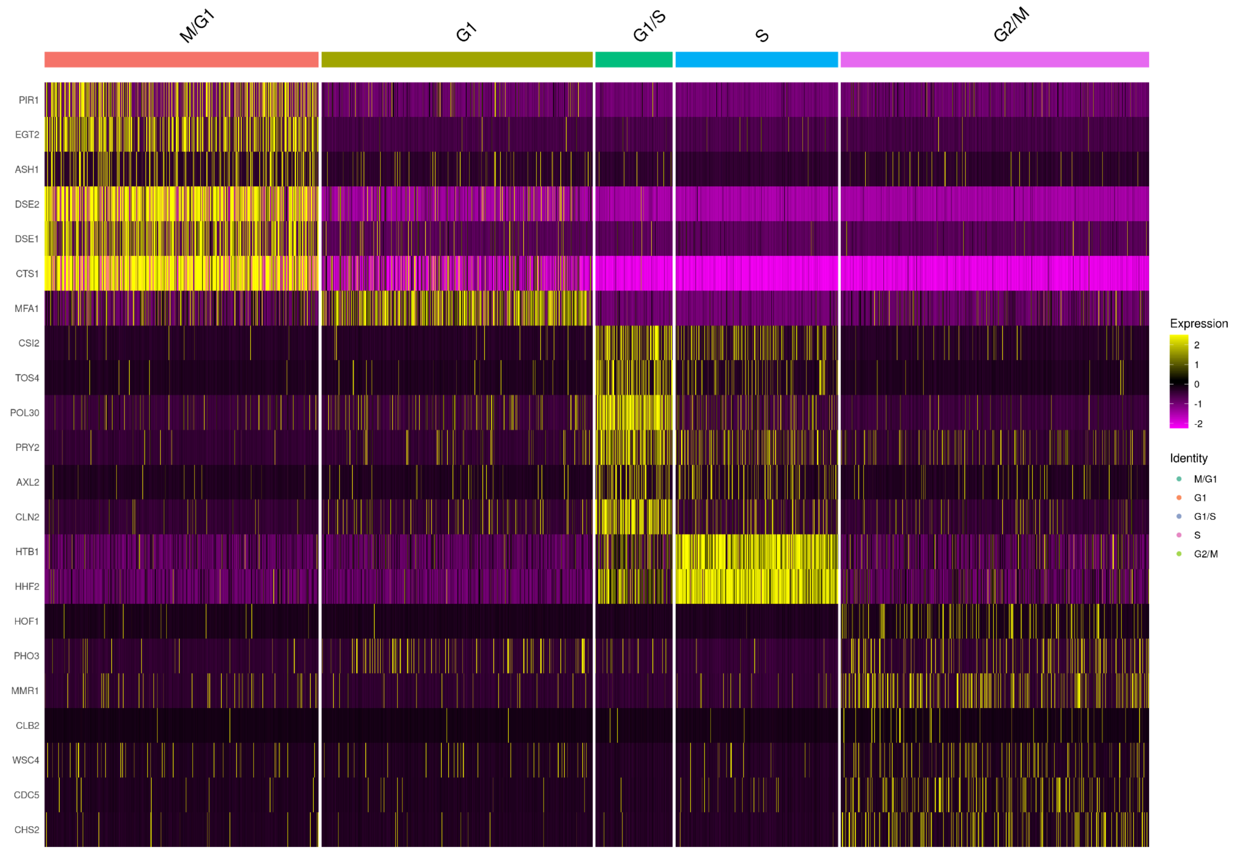


**Figure S3: Marker gene expression of cell-cycle classified single-cells from the set of 393 previously genotyped segregants.** These data are from the first 10x run of the 393 segregants (Number of cells=3,454). The results look similar for the second run. The heatmap shows the normalized expression of each of the 22 markers used for assigning clusters to each cell-cycle stage. For the M/G1 stage, we used the genes *PIR1*, *EGT2*, *ASH1*, *DSE1*, *DSE2*, and *CTS1*. For the G1 stage, we used the gene *MFA1*, which in our experiments reproducibly connected the M/G1 and G1/S transition stages. For the G1/S stage, we used the genes *CSI1*, *TOS4*, *POL30*, *PRY2*, *AXL2*, and *CLN2*. For the S stage, we used these genes *HTB1* and *HHF2*. Finally for the G2/M stage we used the genes *HOF1*, *PHO3*, *MMR1*, *CLB2*, *WSC4*, *CDC5*, and *CHS2*.

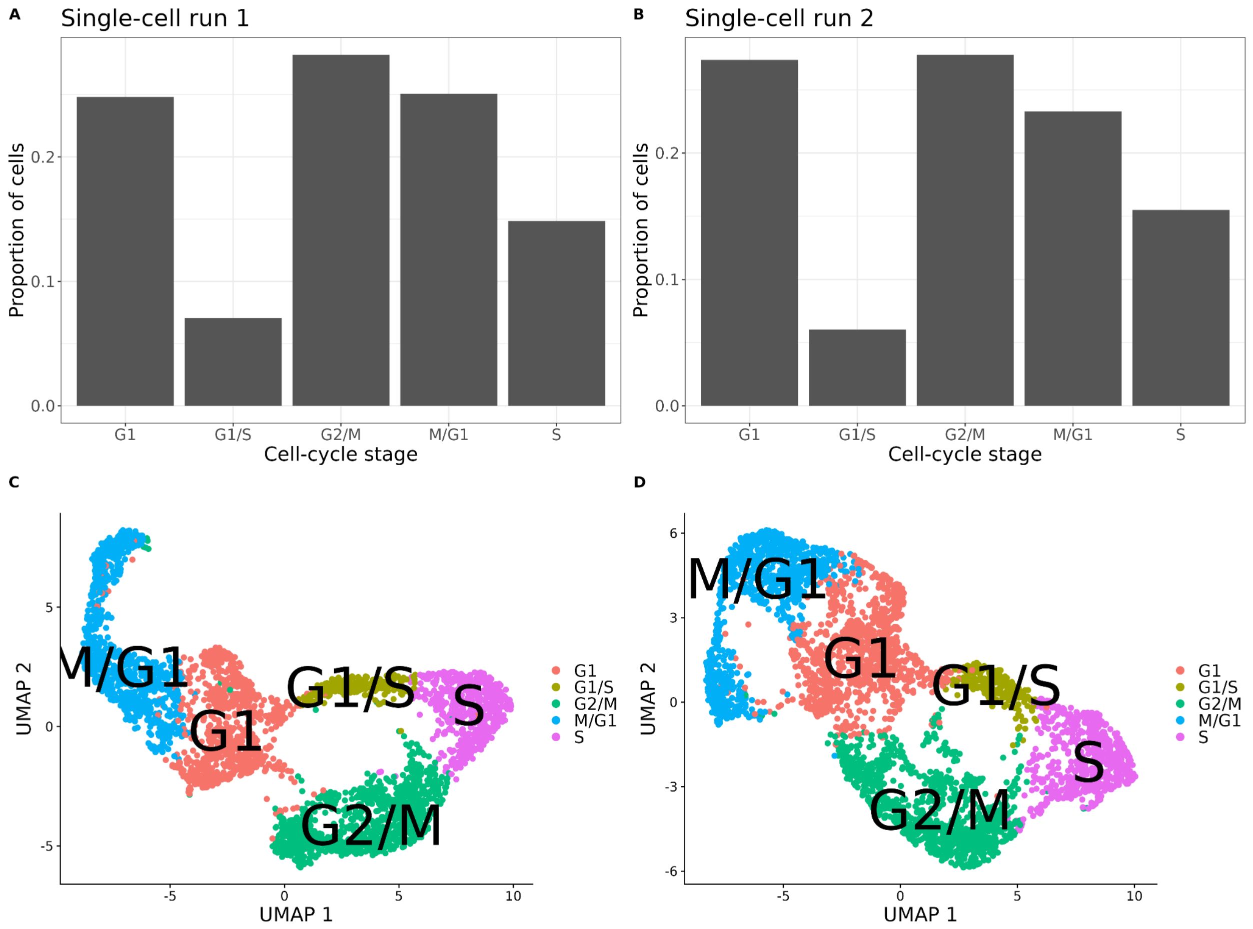


**Figure S4: Cell-cycle classification of the single-cells from the set of 393 previously genotyped segregants. A)** Proportion of cells assigned to each cell-cycle stage for the first of two 10x runs of single-cells from the pool of 393 segregants (Number of cells=3,454). **B)** Proportion of cells assigned to each cell-cycle for the second 10x run of single-cells from the pool of 393 segregants (Number of cells=3,708). **C)** UMAP plot from the single-cell data in A) with cells colored according to their assigned cell-cycle stage. **D)** UMAP plot from the single-cell data in B) with cells colored according to their assigned cell-cycle stage.

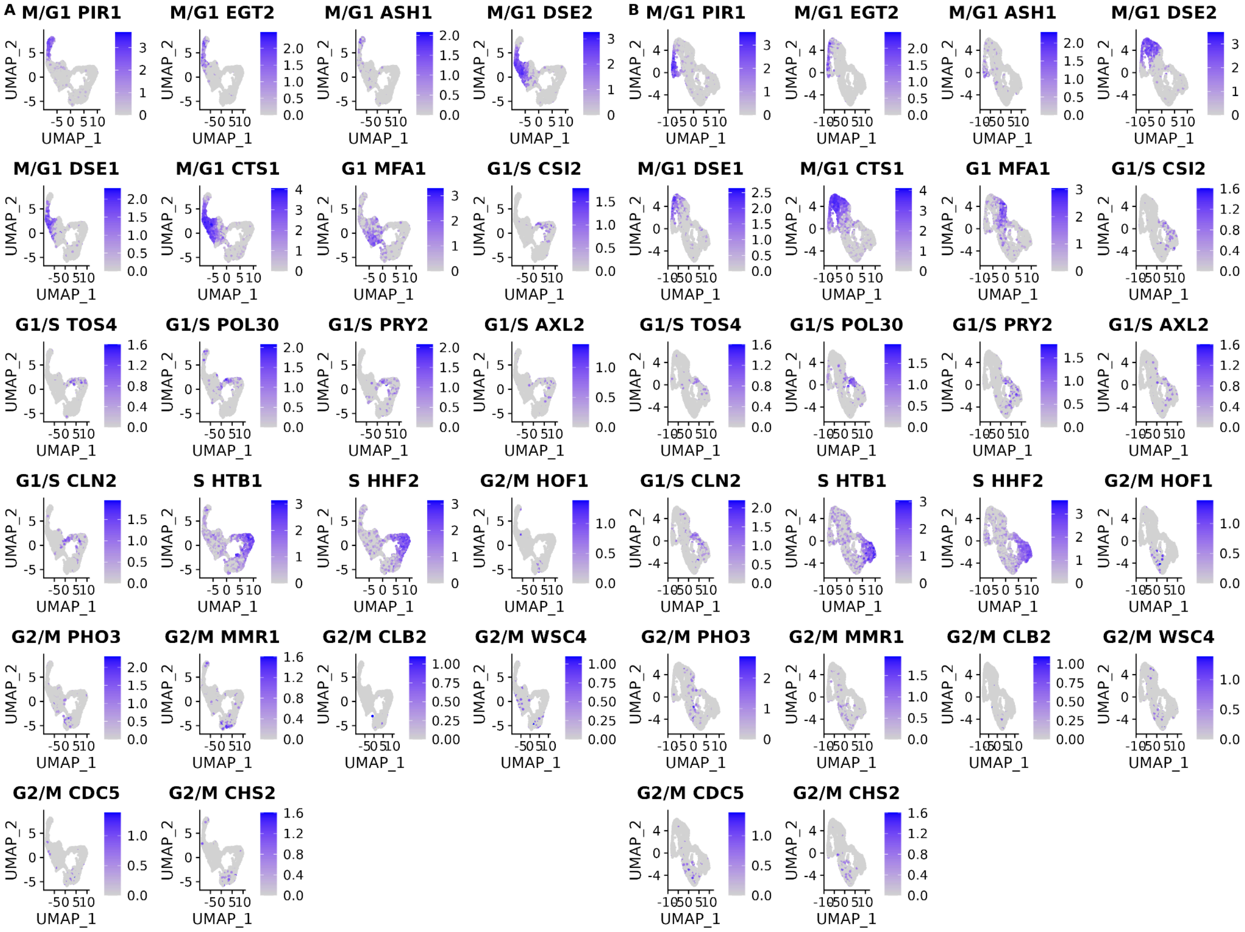


**Figure S5: Cell-cycle marker gene expression of the single-cells from the set of 393 previously genotyped segregants. A)** Gene expression levels of the 18 cell-cycle markers used for classification with panels ordered by their position in the cell-cycle for the first of two 10x runs of single-cells from the pool of 393 segregants (Number of cells=3,454). **B)** same as A) for the second 10x run of single-cells from the pool of 393 segregants (Number of cells=3,708).

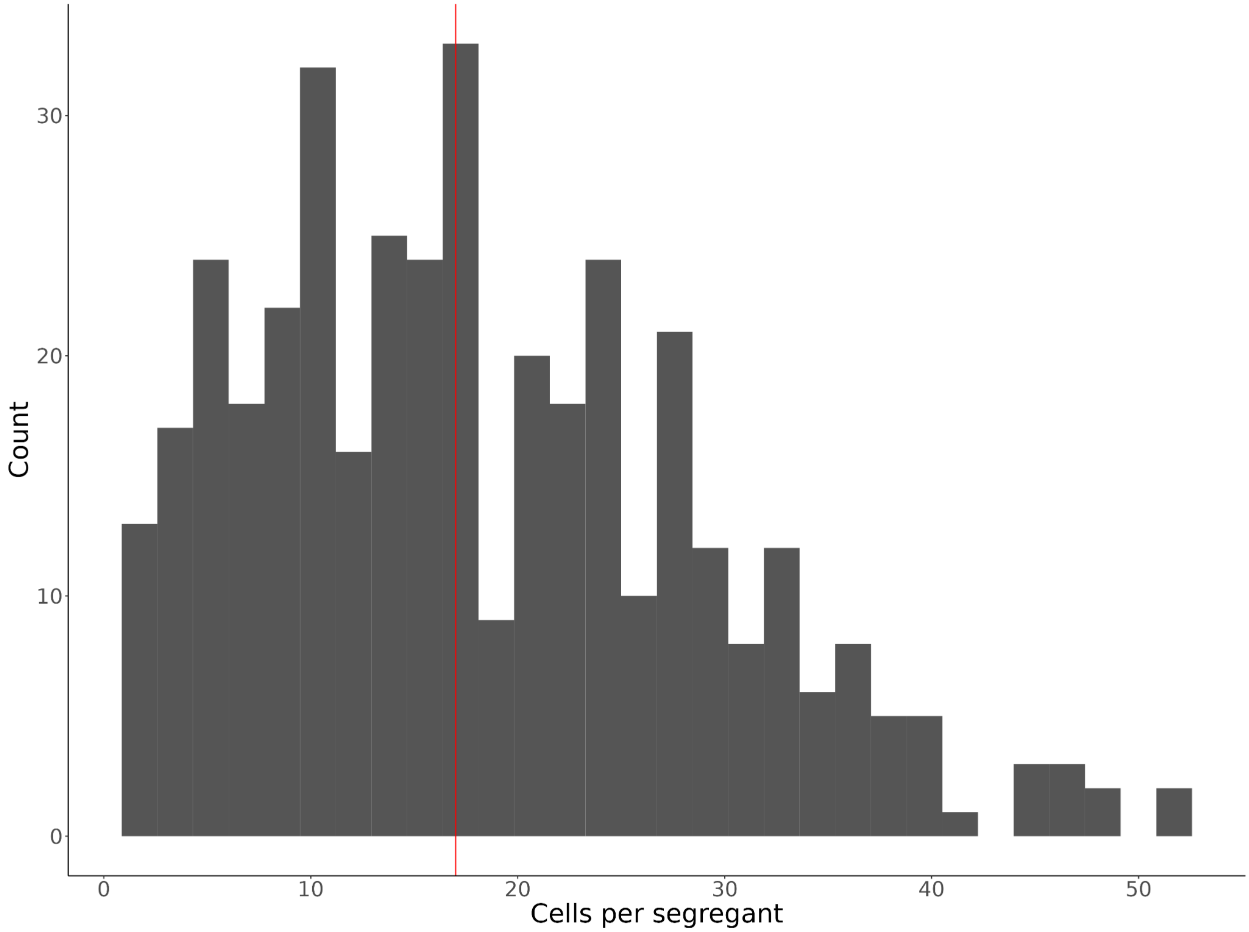


**Figure S6: Histogram of the number of single cells identified for each of the 393 segregants.** The median number of cells per segregant (17 cells) is displayed as a vertical red line.

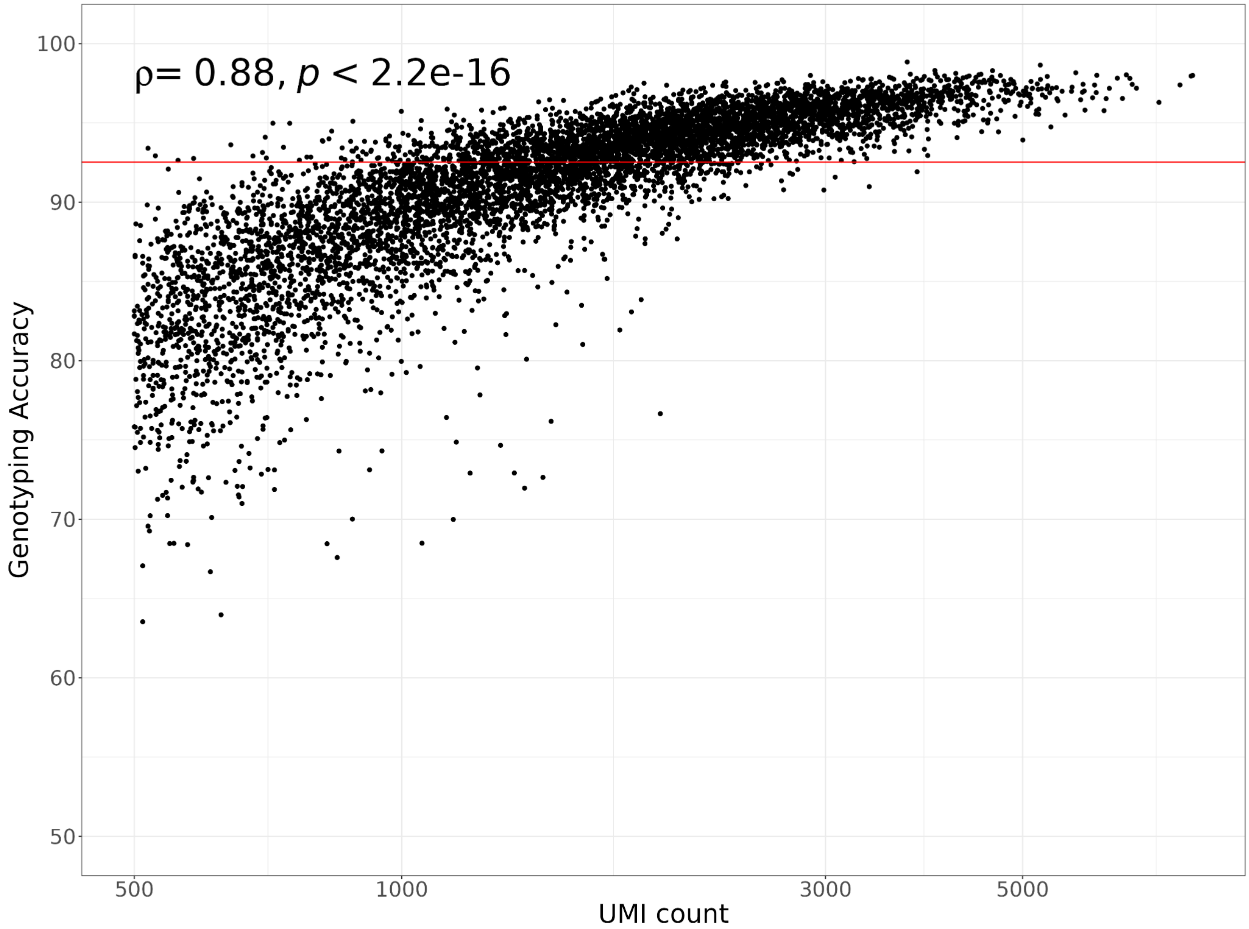


**Figure S7: Single-cell hidden Markov model (HMM) genotyping accuracy compared to the number of unique molecular identifiers (UMIs) per cell.** The horizontal red line shows the median genotyping accuracy of 92.5%  The Spearman's correlation coefficient and p-value comparing the genotyping accuracy to the number of UMIs is shown in the top left corner of the plot. A $\log_{10}$ transformation was applied to the x-axis.

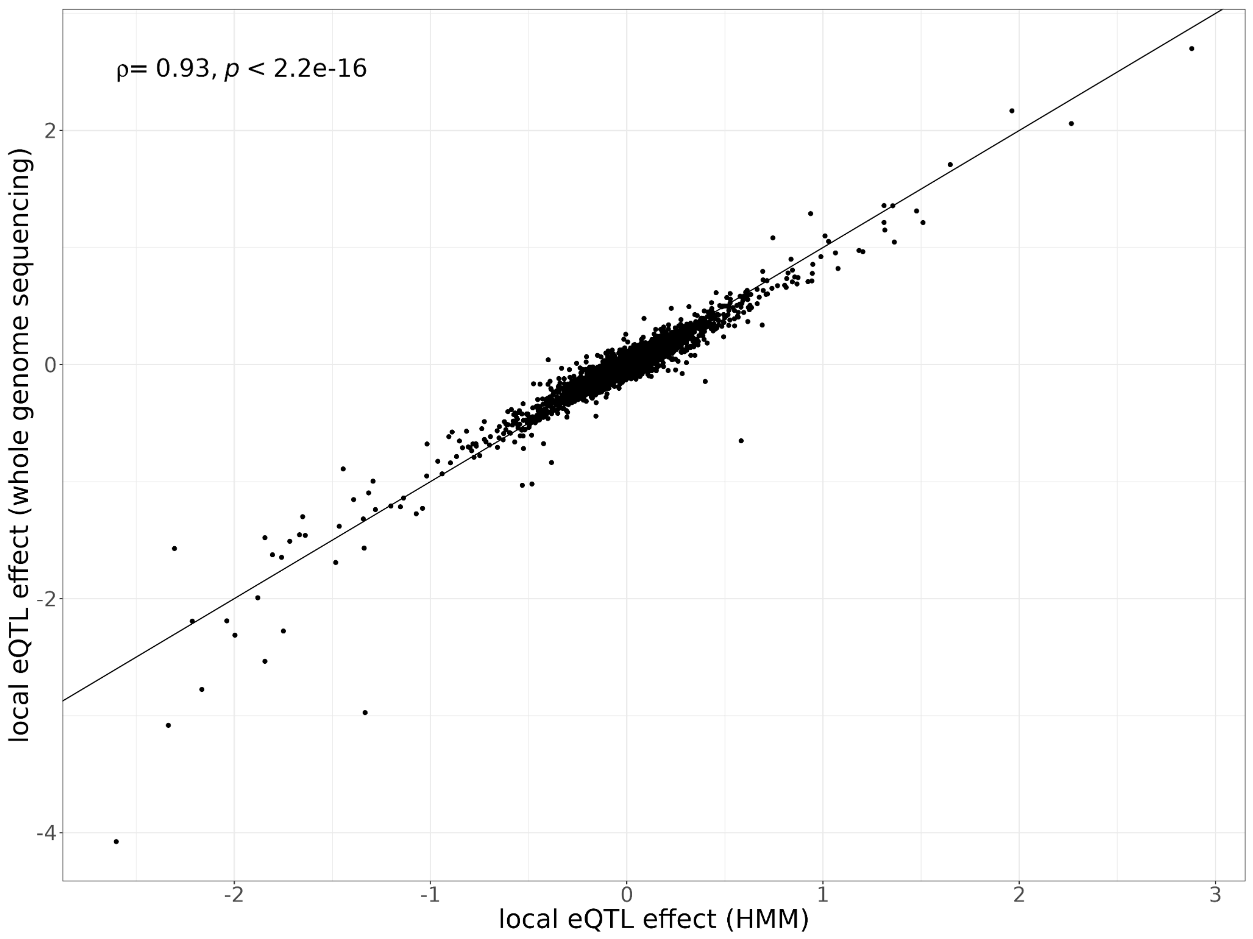


**Figure S8: Local eQTL effects estimated with different genotyping methods.** Local eQTL effect sizes estimated using the HMM-based genotypes from single-cell data (x-axis) compared to local eQTL effect sizes estimated using the lookup of genotypes obtained from whole-genome sequencing (y-axis). The single-cell sequencing data of the 393 segregants is used for quantifying gene expression for each eQTL analysis. The Spearman's correlation coefficient of the local eQTL effect size estimates between these genotyping methods and p-value are shown in the top left corner of the plot.

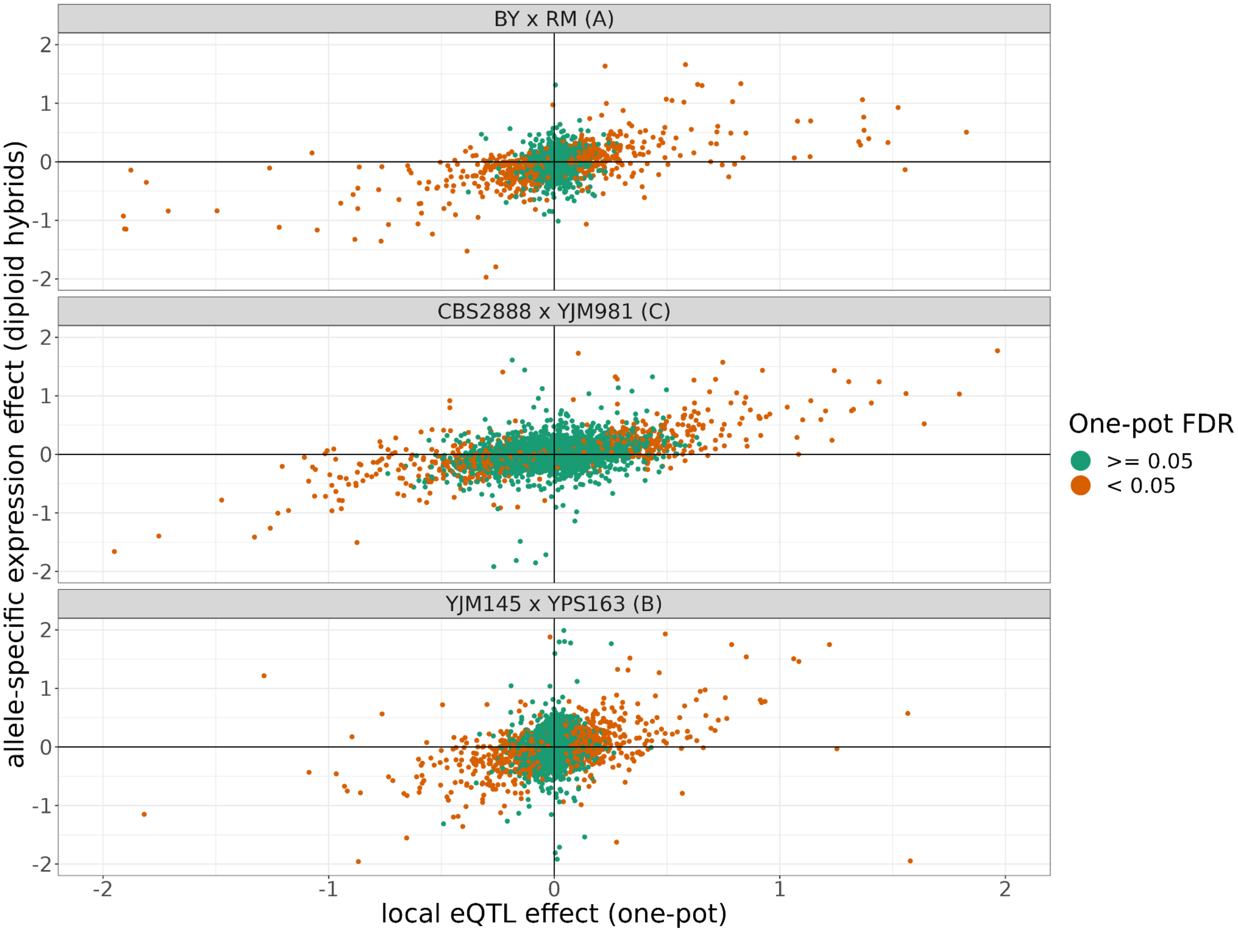


**Figure S9: The allele-specific expression effects compared to the local eQTL effects from each cross.** The position of points on the y-axis represents the allele-specific expression effect of a transcript estimated from the three F1 diploid hybrids of crosses of each of the shown haploid parents and the position of points on the x-axis represents the local eQTL effect estimated for that gene estimated in our one-pot eQTL experiment. The points are colored depending on whether they were significant in the one-pot eQTL experiment at a FDR of <5%. The x and y axes have been truncated at -2 and 2 for ease of visualization purposes, which left out 117 of 7,942 data points.

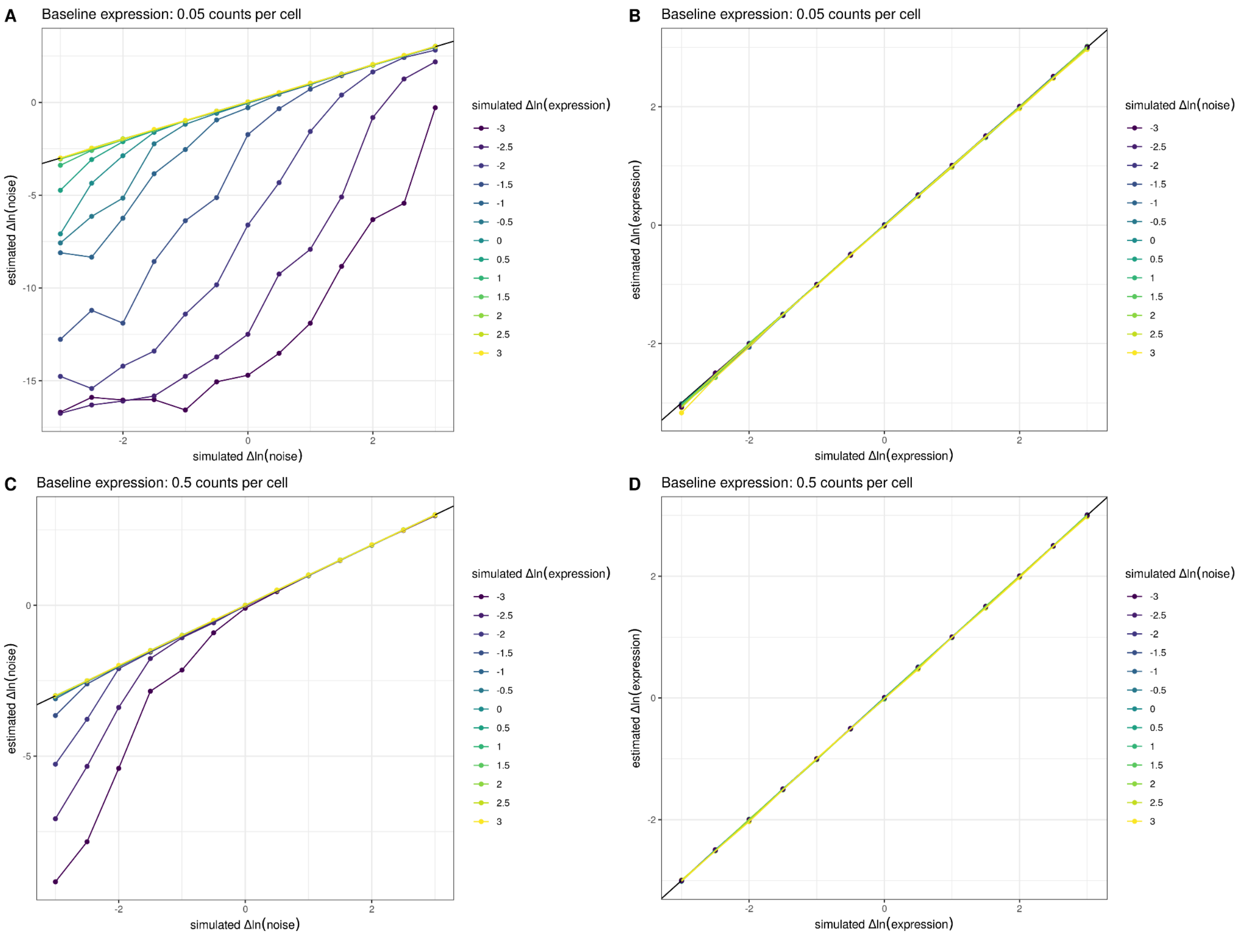


**Figure S10: Estimates of noise are biased downwards at low expression levels.** We simulated allele-specific counts from 5,000 cells across a range of differences in expression level and noise and fit a count-based negative binomial model with a genotype effect for both the mean and the noise (Methods). The estimated and simulated parameters are displayed across the panels. For **A)** and **B),** we assumed a baseline expression of 0.05 counts per cell, and for **C)** and **D),** we assumed a baseline expression of 0.5 counts per cell. 0.05 counts per cell represents the number of counts observed for an average yeast gene in our experiments, and 0.5 counts per cell represents the average number of counts observed for the top 10% of yeast genes in our single-cell data. In **A)** and **C)** the relationship between the simulated change in ln(noise) between alleles and estimated change in ln(noise) between alleles is shown. Individual lines were grouped and colored based on the amount of simulated change in ln(expression) between alleles. In **B)** and **D)**, the relationship between the simulated change in ln(expression) between alleles and estimated change in ln(expression) between alleles is shown. Individual lines were grouped and colored based on the amount of simulated change in ln(noise). These simulations show that the estimates of noise are biased downwards when expression levels are low, but the p-values are well calibrated and not significant in such instances (data not shown). This behavior of the models runs opposite to the global trend we observe whereby increasing expression decreases noise.

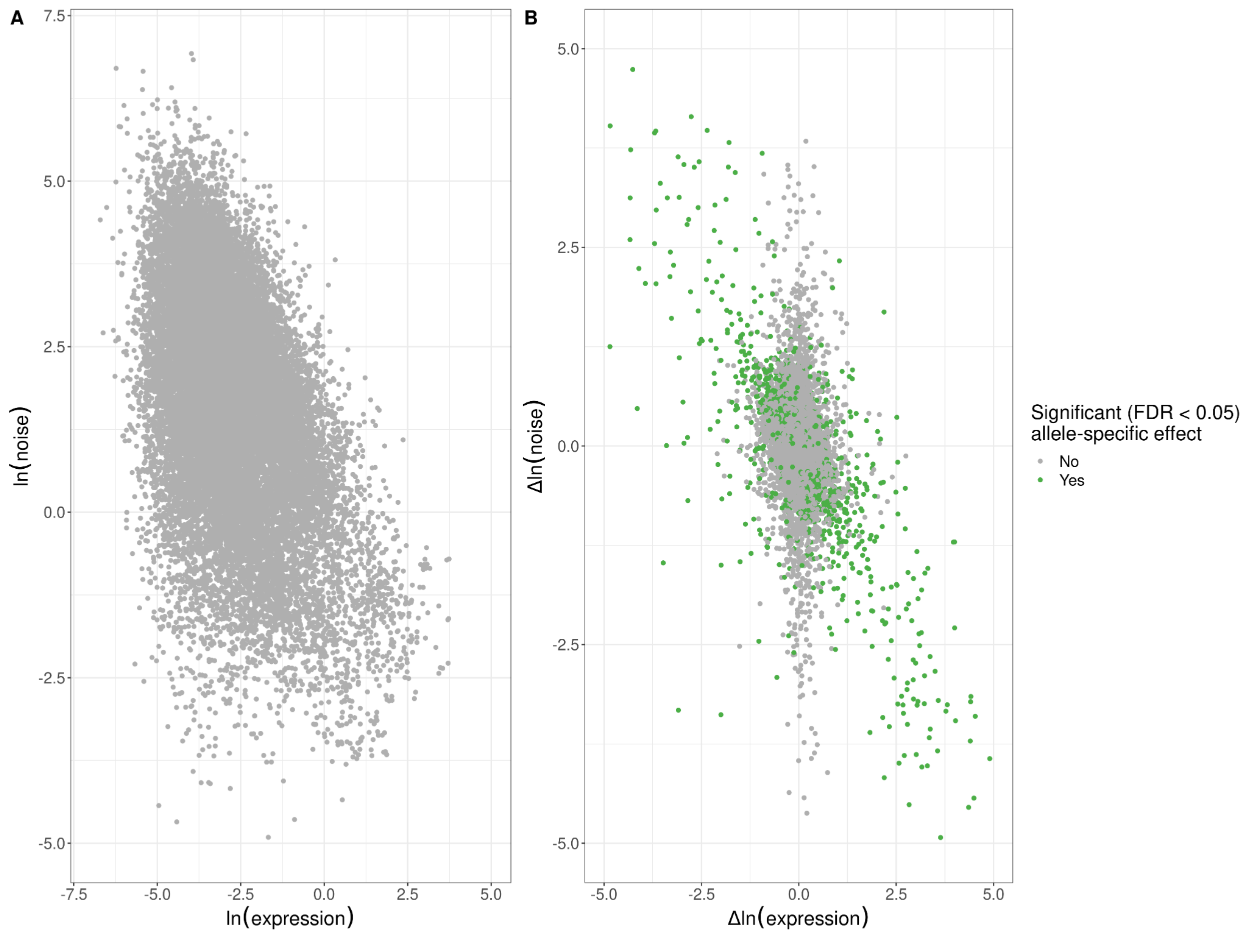


**Figure S11: Empirical global relationship between expression noise and average expression levels. A)** Log-log scatter plot of allelic average expression (x-axis) against allelic expression noise (y-axis); points correspond to all 22,294 alleles for which we were able to estimate allele-specific effects. Average allelic expression is negatively correlated with allelic noise (Spearman's $\rho$=-0.42, p<$10^{-15}$). **B)** Log-log scatter plot of change in expression between alleles (x-axis) against change in expression noise between alleles (y-axis); points correspond to all 11,147 genes for which we were able to estimate allele-specific effects. Points in green highlight the 1,487 genes with significant allele-specific effects at an FDR of <5% on expression noise and/or average expression. Changes in noise are negatively correlated with expression level, across all genes (Spearman's $\rho$=-0.32, p<$10^{-15}$), genes without an allele-specific effect (Spearman's $\rho$=-0.23, p<$10^{-15}$), and genes with significant allele-specific effects (Spearman's $\rho$=-0.64, p<$10^{-15}$). The x and y axes have been truncated at -5 and 5 for ease of visualization purposes, which left out 142 of 11,147 data points.

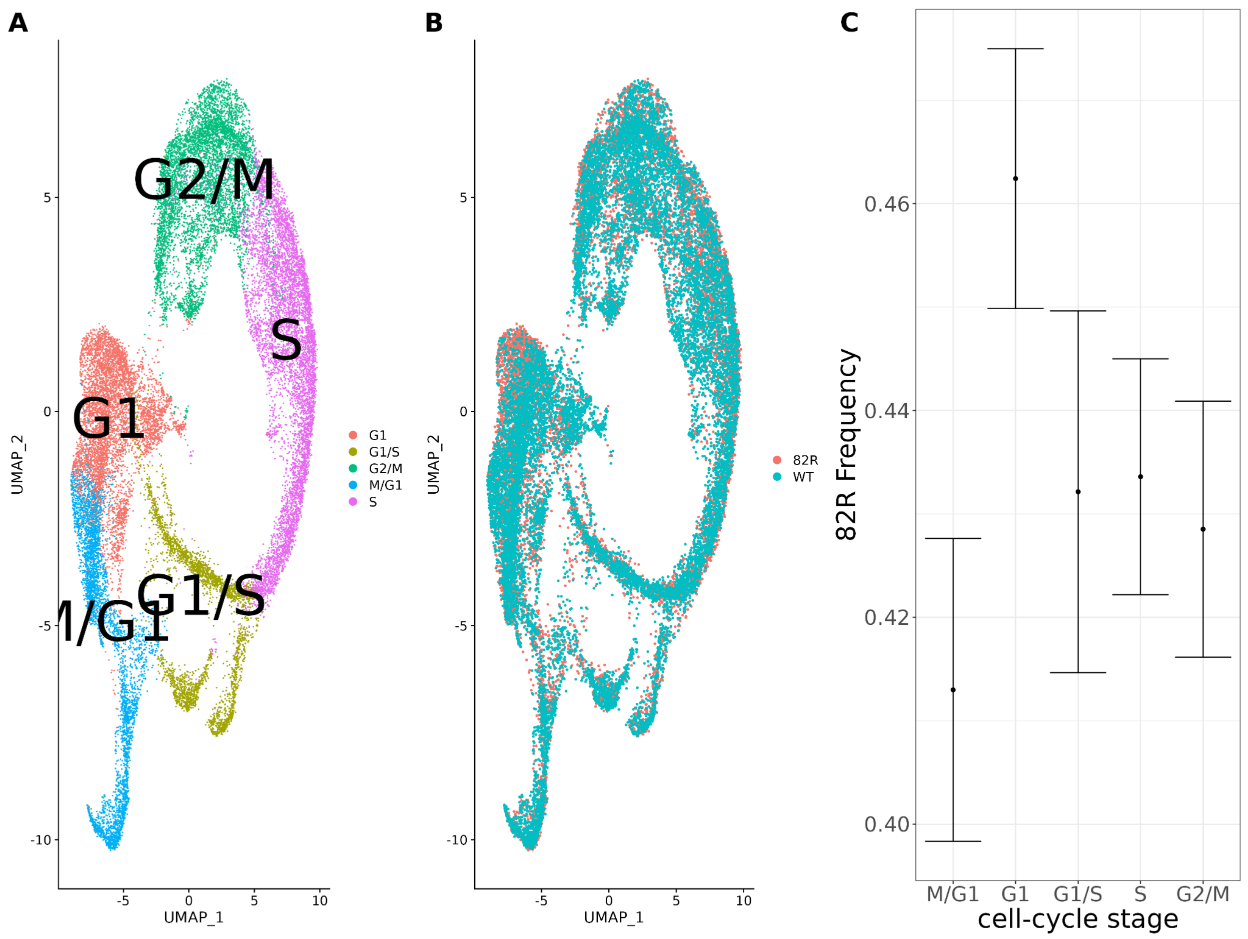


**Figure S12: Single-cell expression profiling of allele-replacement strains comparing the cell-cycle distribution of strains with the 82R allele of *GPA1* to strains with the WT allele of *GPA1*.** scRNA-seq of 26,859 cells from 11,695 cells with the 82R allele and 11,695 cells with the WT (82W) allele. **A)** Combined UMAP plot of the integrated single-cell dataset with cells colored according to their cell-cycle stage. **B)** Combined UMAP plot of the integrated single-cell dataset with cells colored according to their genotype at position 82 of the GPA1 protein. **C)** Allele-frequency of the 82R allele across the cell-cycle in our allele-replacement single-cell data. Error bars represent the 95% confidence intervals for the proportion of cells in each cell-cycle stage.

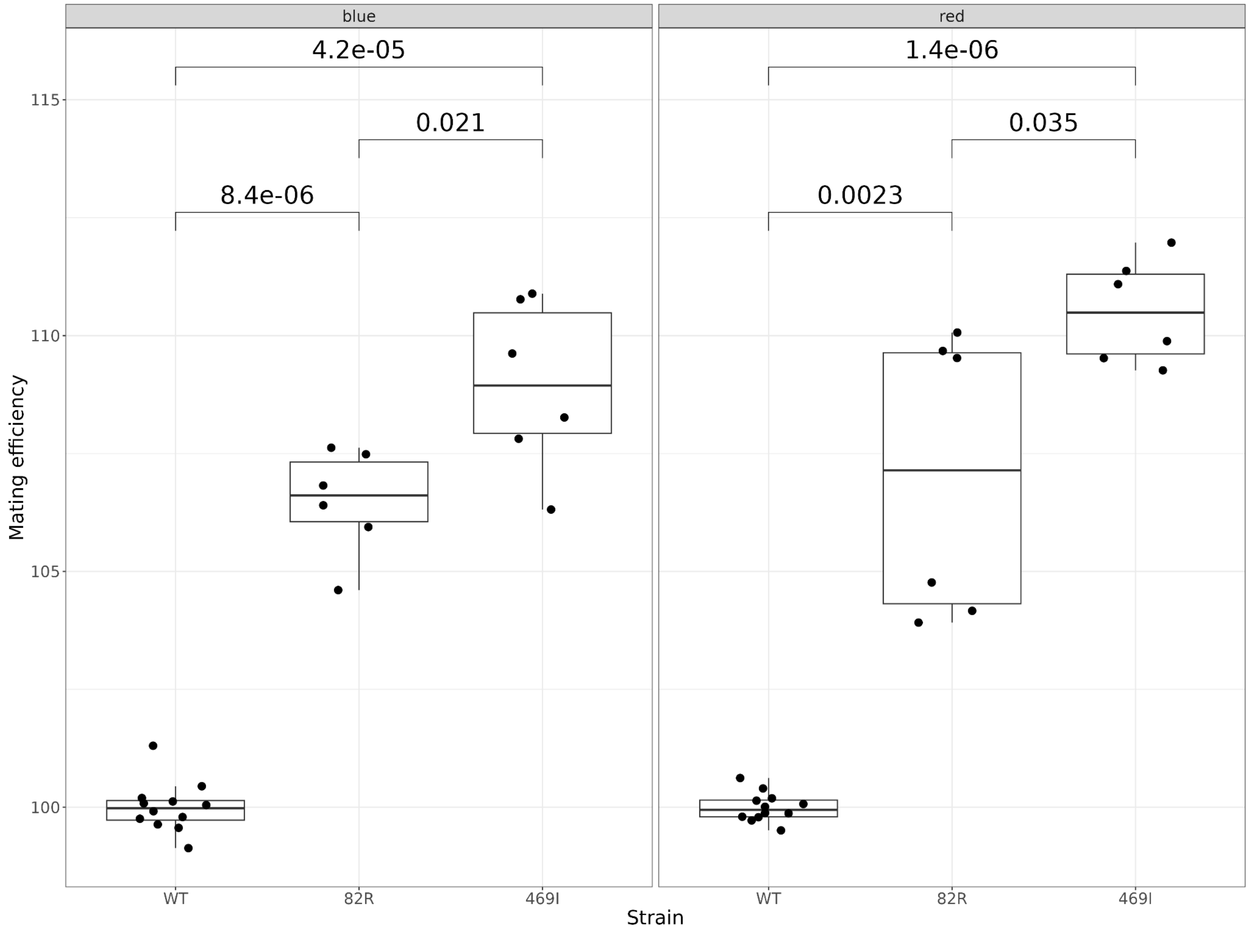


**Figure S13: Mating efficiency of the 82R allele of *GPA1* compared to the 469I and wild-type (82W 469S) alleles.** The boxplots show the mating efficiency of different allele-replacement strains. Each point represents replicate measurements of the mating efficiency as estimated by flow cytometry. The plot is further subdivided into two panels depending on the color of the fluorescent protein used to estimate mating efficiency. Mating efficiency values were normalized to the average mating efficiency value of the WT strain. The unadjusted p-values of pairwise t-tests between the alleles are shown.

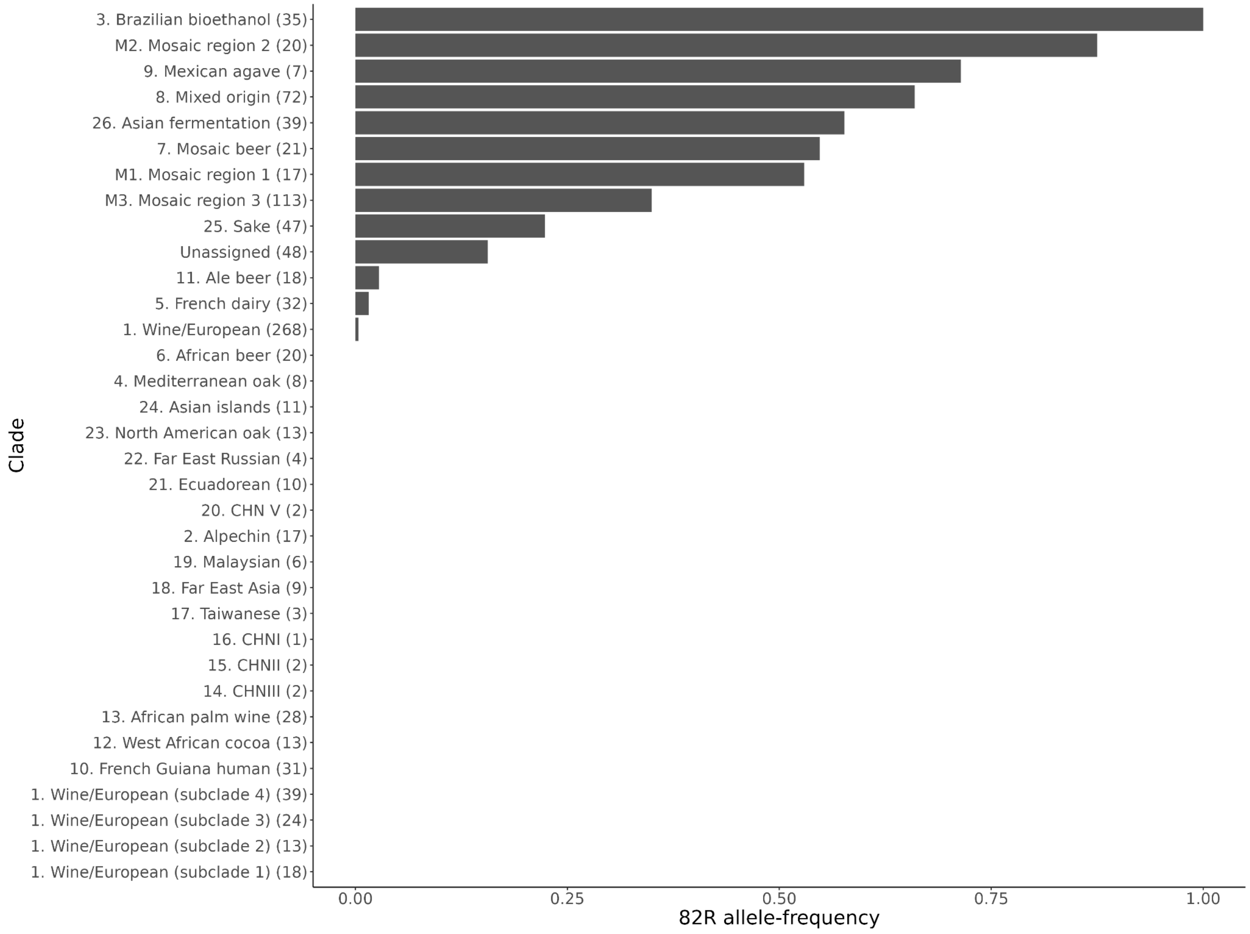


**Figure S14: Allele-frequency of the 82R allele of *GPA1* in sequenced yeast strains.** The allele-frequency is further broken down by the clade assignments given in Peter et al.[39]. The number of strains in each clade is given in the brackets next to the clade name.

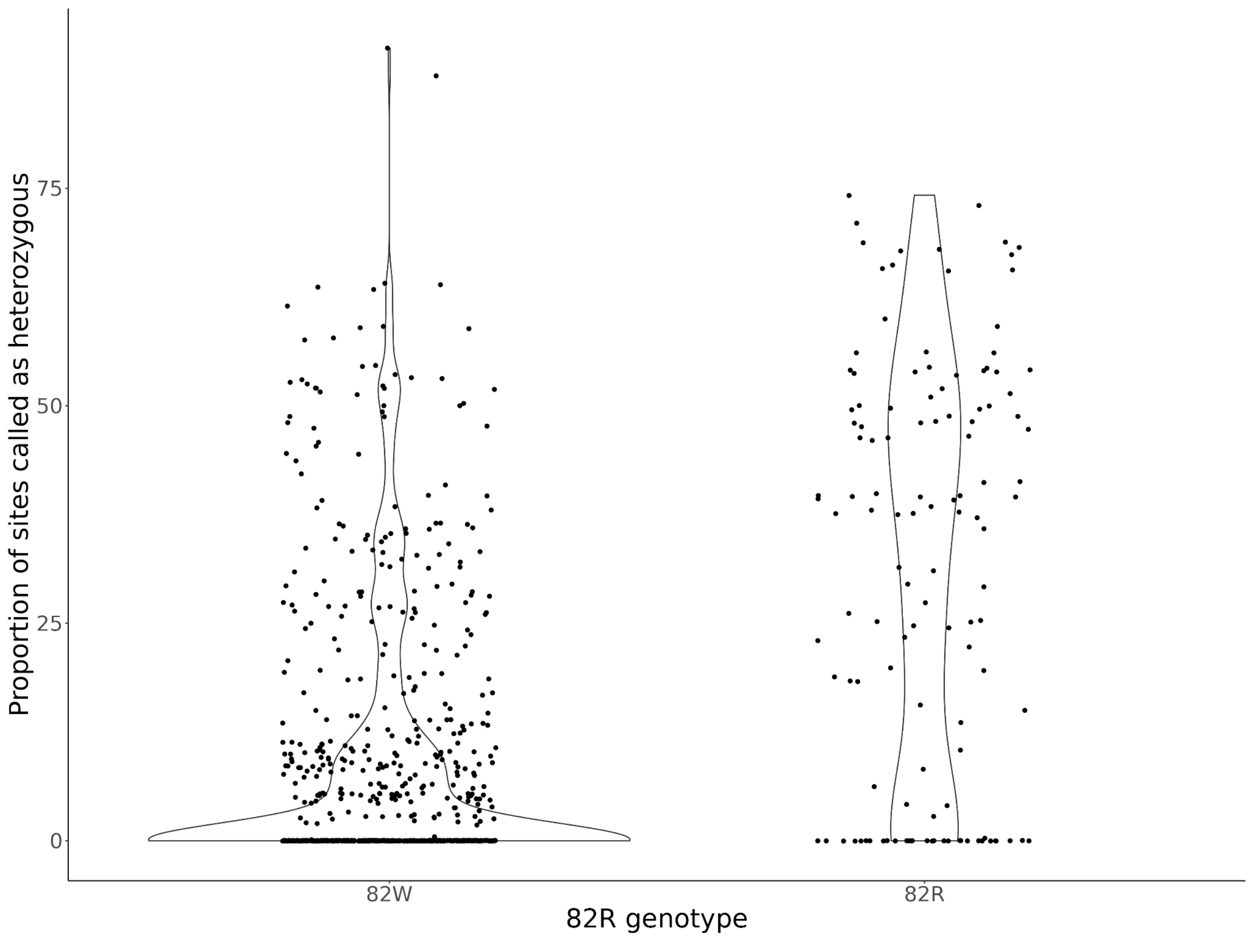


**Figure S15: Genome-wide heterozygosity per strain, after classifying strains as having the 82W allele or 82R allele of *GPA1*.** The violin plots display the distribution of the proportion of sites per strain called heterozygous from Peter et al.[39]. Strains were classified depending on whether it is homozygous for the 82W (N=760) or 82R (N=164) alleles. Strains with support for both 82R and 82W alleles (N=87) were not analyzed here.

**Supplementary Tables**

**Table S1:** Strains used in this study.

**Table S2:** Plasmids used in this study.

**Table S3:** Primers used in this study.

**Table S4:** Summary information for the single-cell expression data generated in this study. For the ASE data, the number of transcripts per cell refers to the number of transcripts with transcribed variants.

**Table S5:** Variance explained by cell-cycle stage and the effect of segregant per transcript for the 393 previously generated segregants. The statistical significance of the segregant and cell-cycle stage effects for each transcript was estimated using a likelihood ratio test, and a corrected p-value is reported. Missing q-values indicate that the model did not converge for that gene.

**Table S6:** Local eQTL summary statistics for the 393 previously generated segregants from the BY and RM cross. Effect size (Betas) and p-values with the 'old' suffix were estimated using the genotypes from whole-genome sequencing. Betas and p-value with the 'hmm' suffix were estimated using the genotypes inferred using our HMM.

**Table S7:** Local eQTL summary statistics for our one-pot eQTL experiments. Each sheet has the local eQTL summary statistics for the three crosses we examined (cross A=BY and RM, cross B=YJM145 and YPS163, cross C=CBS2888xYJM981). The column 'has cell-cycle interaction' is set to 1 if a cell-cycle interaction was observed at a FDR of <5%. For cross A, the summary statistics from bulk eQTL[7] are provided in additional columns. Missing values in the bulk eQTL columns indicate that a gene did not pass our filtering criteria, and a local eQTL test was not performed.

**Table S8:** Distant eQTL summary statistics for our one-pot eQTL experiments. Each sheet has the local eQTL summary statistics for the three crosses we examined (cross A=BY and RM, cross B=YJM145 and YPS163, cross C=CBS2888xYJM981). The column 'has cell-cycle interaction' is set to 1 if a cell-cycle interaction was observed at a FDR of <5%. The column 'eQTL in hotspot' is set to 1 if a distal eQTL falls within a significant hotspot bin.

**Table S9:** Single-cell allele-specific expression (ASE) summary statistics. Each sheet has the ASE summary statistics for the parental hybrids of the three crosses we examined (cross A=BY and RM, cross B=YJM145 and YPS163, cross C=CBS2888xYJM981). The local eQTL summary statistics were included from the corresponding one-pot eQTL experiment for each transcript that was tested there. Missing values in the one-pot eQTL columns indicate that a gene did not pass our filtering criteria, and a local eQTL test was not performed.

**Table S10:** Summary statistics for allele-specific effects on noise and average expression. Each sheet has the summary statistics for the parental hybrids of the three crosses we examined (cross A=BY and RM, cross B=YJM145 and YPS163, cross C=CBS2888xYJM981). The estimates and p-values are derived from the joint model that contains cell-cycle, cell-cycle interactions, and allelic effects on both the mean and noise. The column 'Overlaps global trend line' is set to 1 if the 95% confidence interval of the noise effect did not overlap the 95% confidence interval of the global trend line.

**Table S11:** Cell-cycle occupancy QTL summary statistics for our one-pot eQTL experiments. Each sheet has the cell-cycle occupancy QTL summary statistics for the three crosses we examined (cross A=BY and RM, cross B=YJM145 and YPS163, cross C=CBS2888xYJM981).

**Table S12:** Single-cell RNA sequencing of allele-replacement strains with the 82R and 82W alleles of *GPA1.* The distant eQTL effects near *GPA1* from our one-pot eQTL experiment are contrasted to a differential expression analysis of our allele replacement strains. The p-values from our one-pot eQTL study were adjusted using a permutation procedure, and a Bonferroni correction was used to adjust the single-cell validation p-values. Missing values in the single-cell validation columns indicate that a gene did not pass our filtering criteria, and a differential expression test was not performed.

**File S1:** Hotspot annotation files for each cross (cross A=BY and RM, cross B=YJM145 and YPS163, cross C=CBS2888xYJM981, A_bulk=BY and RM with expression measured with bulk RNA-seq[7]).

**File S2:** Alignment and tree files from a clustered blast search of the yeast Gpa1 protein sequence (YHR005C).
