## Supplementary material for "Single-cell eQTL mapping in yeast reveals a tradeoff between growth and reproduction": File S1: README.md

## Annotation spreadsheets for all the distal trans eQTL hotspots identified in this study.

### Data for "Single-cell eQTL mapping in yeast reveals a tradeoff between growth and reproduction"


The sub-folders collate the result from the the three crosses we examined using single-cell eQTL mapping (A, B, C), and our reanalysis of Albert et al 2018 (A_2018)

Each folder contains an annotation spreadsheet for every distal trans eQTL hotspot identified. The files are named according to the hotspot window and number of markers.
The naming scheme is as follows `{chr}:{left}-{right}_{genecount}_finemapping.xlsx` 

### There are 13 different sheets in each spreadsheet, a description of these sheets can be found below. 

#### **cell_cycle_causals**

Lists the cell-cycle occupancy QTL that overlap the hotspot CI. Data will be NONE if there is no overlap

#### **cell_cycle_assoc**

Lists the cell-cycle association at the peak marker of the most significantly associated transcript in the hotspot. 

#### **causals_round_robin**

Lists causal genes that overlap the hotspot CI from the same cross using the round robin data from Bloom et al. 2019. Data will be NONE if there is no overlap.

#### **assoc_round_robin**

List any QTL identified in Bloom et a 2019 that overlap the hotspot window from the same cross. Data will be NONE if there is no overlap.

#### **complete_assoc**

Association statistics for 40 different quantitative traits at the peak marker of the most significantly associated transcript in the hotspot.

#### **provean_snps**

Annotated SNPs within each hotspot CI. The allele frequency in the 1000 yeast genomes collection and the PROVEAN score is provided if available. 

#### **GO**

Significant Gene Ontology (GO) term enrichments for the genes linked to each hotspot.

#### **KEGG**

Significantly Kyoto Encyclopedia of Genes and Genomes (KEGG) term enrichments for genes linked to each hotspot.

#### **GO_raw**

Full GO term enrichment table. 

#### **Directional hotspot distal**

List of all distant eQTLs for the hotspot. Columns include effect sizes for each cell-cycle (BETA_{stage}) and overall (Beta,SE), the CI of the eQTL, and whether the eQTL has an interaction with the cell-cycle (has_interaction).

#### **Directional hotspot local**

List of all local eQTLs with the hotspot CI. Columns include effect sizes for each cell-cycle (BETA_{stage}) and overall (Beta,SE), the CI of the eQTL, and whether the eQTL has an interaction with the cell cycle (has_interaction).

#### **genes_in_region**

List of all genes in the hotspot CI.

#### **eqtl_region**

Hotspot CI: chrom, start, end


 
